# Computational design of a multi-epitope vaccine against M. tuberculosis

**DOI:** 10.64898/2026.07.09.737463

**Authors:** Abdulrasheed Buhari, Okutu Peter, Adithya Sivakumar, Oyeleke Usman Abolaji, Hameed Sodiq Ayobami

## Abstract

**Background:** Tuberculosis remains a leading global infectious killer, with BCG offering inconsistent adult protection and rising drug-resistant strains demanding novel vaccine strategies. We report the first multi-epitope vaccine construct simultaneously targeting three previously unexplored *Mycobacterium tuberculosis* virulence proteins; EccB3, MycP, and polyketide synthase which collectively govern nutrient acquisition, ESX secretion integrity, and innate immune evasion.

**Methods:** Using a reverse vaccinology pipeline, B-cell, CTL, and HTL epitopes were predicted, filtered for allergenicity, toxicity, and IFN-γ induction, then assembled into an 823-residue chimeric construct incorporating beta-defensin and PADRE adjuvants with AAY/GPGPG linkers, covering ∼90% global HLA diversity. The construct underwent AlphaFold structure prediction, 3DRefine refinement, disulfide engineering, PROCHECK/ProSA validation, ClusPro 2.0 docking against TLR1/TLR2, and C-IMMSIM immune simulation.

**Results:** The construct (82.3 kDa, instability index 32.48) showed strong structural quality (94.7% favoured Ramachandran residues), stable TLR1/TLR2 binding (weighted energy: −1,371.0 kcal/mol), and robust in silico immune responses and durable memory cell formation following booster simulation.

**Conclusion:** This computationally validated construct represents a promising multi-target TB vaccine candidate warranting experimental advancement.

## 1.0 INTRODUCTION

Tuberculosis is caused by *Mycobacterium tuberculosis* (Mtb) and remains a major global health burden responsible for more than 1.3 million deaths annually. It is estimated that about one fourth of the global population is latently infected with Mtb[1]. The only licensed vaccine, Bacillus Calmette-Guérin (BCG) offers variable protection against pulmonary TB in adults, and growing prevalence of multidrug-resistant (MDR) and extensively drug resistant (XDR) Mtb strains has necessitated the need for better preventive measures[2]. The key to the challenges of Mtb’s is their ability to suppress host macrophage responses using specialized secretion systems, surface lipid modifications and the capability to actively obtain nutrients even during latent infection phase [3,4]. The use of reverse vaccinology and multi-epitope vaccine design provides a more rational and promising approach compared to traditional methods of designing and developing vaccines against the bacteria. Bioinformatic prediction of immunogenic targets from the pathogen’s proteome and the use of selected epitopes to design a chimeric structure as a vaccine candidate allows for simultaneous activation of cytotoxic T lymphocytes (CTLs), helper T lymphocytes (HTLs) and B cells [5,6]. For Mtb, vaccination would need to induce the stimulation and activation of IFN-γ-secreting CD4+ and CD8+ T cells and engagement of pattern recognition receptors such as TLR1/TLR2 heterodimer through which mycobacterial lipoproteins and cell wall components are sensed by antigen presenting cells like macrophages and dendritic cells [7].

Although immunoinformatics approaches have been used in other studies aimed at computationally designing vaccines against Mtb, the three targeted proteins used in this study, EccB3, MycP, and polyketide synthase have not previously been investigated individually or in combination for epitope profiling or multi-epitope vaccine construction. EccB3 is a membrane-associated component of the ESX-3 type VII secretion system and is essential for iron and zinc acquisition from the host macrophage environment. This function is critical for the viability of intracellular Mtb [8]. MycP is a membrane-anchored mycosin serine protease that plays a major role in the ESX secretion system assembly/stabilization and virulence. It has a periplasmic domain that is exposed and potentially accessible to antibody-mediated inhibition[9]. Polyketide synthase mediates biosynthesis of phthiocerol dimycocerosate (PDIM) and related cell wall lipids that mostly mask pathogen-associated molecular patterns from innate immune recognition and disrupts phagolysosomal membranes to enhance the survival of the bacterial intracellularly[10]. Together, these targets address Mtb’s nutrient acquisition, secretion-mediated virulence, and immune evasion.

In this study (with the overview described in figure 1), epitopes predicted from these three proteins were assembled into a chimeric construct incorporating beta-defensin and PADRE as adjuvant sequences, with AAY and GPGPG linkers, using HLA alleles covering approximately 90% of the global population. The construct was evaluated through AlphaFold tertiary structure prediction, 3D refinement, disulfide bond engineering, ProSA and PROCHECK validation, ClusPro 2.0 molecular docking against the TLR1/TLR2 heterodimer (PDB: 2Z7X), and normal mode analysis of the docked complex via iMODS.

**Figure 1.**
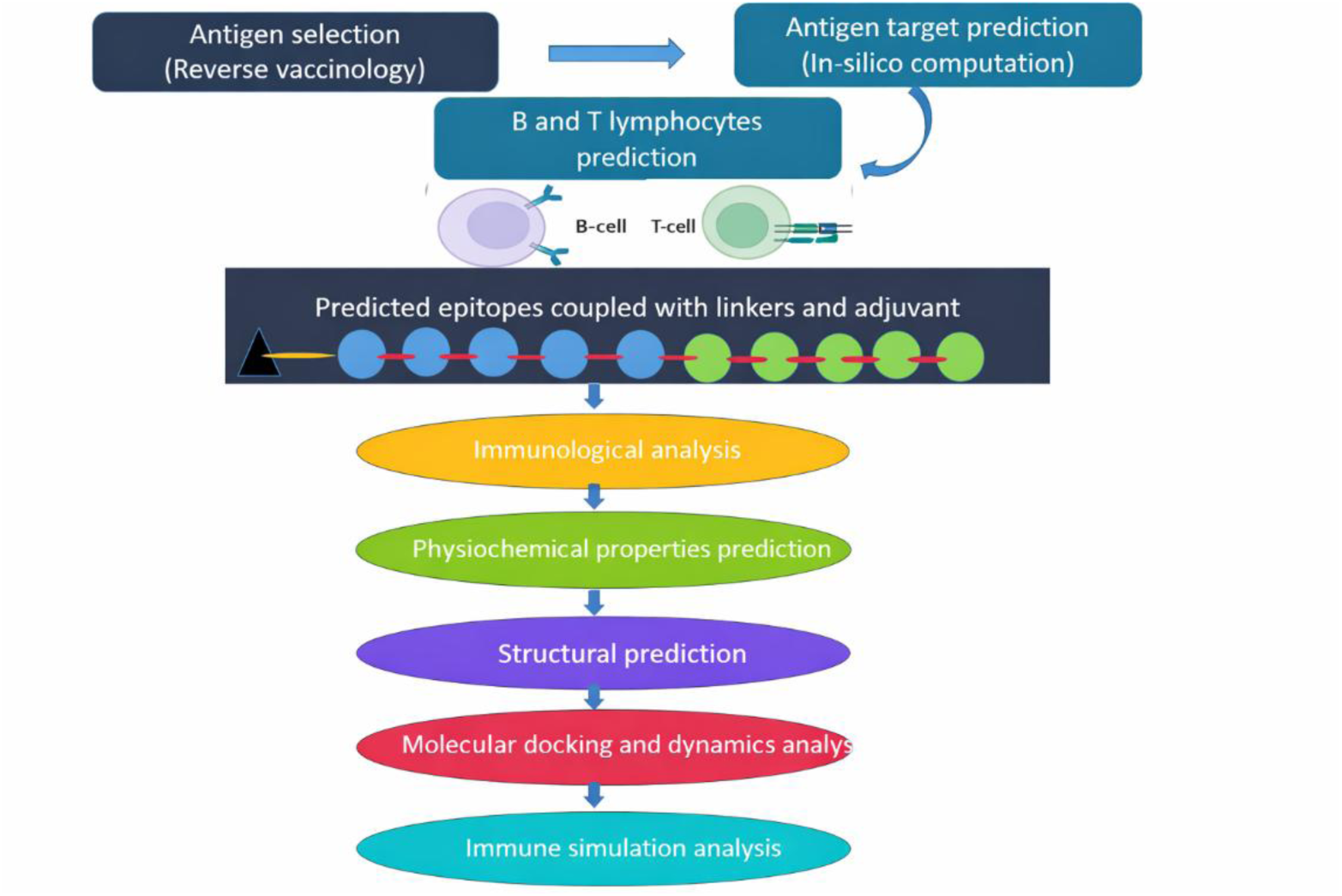
Schematic overview of the computational approach employed in the development of the Mtb multi-epitope vaccine construct

## 2.0 MATERIALS AND METHOD

### 2.1 Antigen Retrieval and Antigenicity Assessment

The amino acid sequences of selected *Mycobacterium tuberculosis* proteins were retrieved from publicly available protein databases and screened for antigenic potential using VaxiJen (http://www.ddg-pharmfac.net/vaxijen/VaxiJen/VaxiJen.html). This is an alignment-independent prediction tool that uses auto cross-covariance (ACC) transformation of amino acid properties to evaluate antigenicity and identify likely antigens, without considering sequence homology. This method allows quick detection of proteins that have inherent immunogenic capability that can be used in designing a vaccine [11].

A bacterial model was used with a threshold value of 0.4 and only proteins with greater than this cut off were regarded as probable antigens. Such a screening process is important to filter out candidates that lack enough physicochemical traits that are linked to immune recognition and this way, there are more chances that epitopes will be presented and the immune system activated effectively[11]. The proteins that passed this requirement were then further progressed in epitope prediction studies, which underlie vaccine design based on downstream immunoinformatics analyses.

### 2.2 Helper and Cytotoxic T Lymphocyte (HTL) Epitopes Prediction

Cytotoxic and helper T lymphocyte (CTL) epitopes were predicted using the NetCTLpan 1.2 server, accessible through the Immune Epitope Database (IEDB). This tool integrates three complementary biological processes to identify candidate MHC class I-restricted peptides (https://tools.iedb.org/mhci/): (i) binding affinity prediction for MHC class I molecules, (ii) proteasomal cleavage efficiency, and (iii) transporter associated with antigen processing (TAP) transport efficiency [12]. A combined threshold score of ≥0.75 was applied, retaining only those peptides predicted to be strong MHC-I binders with efficient intracellular processing. Prediction of MHC class II binding epitopes were done to find potential CD4+ T cell activating regions. Selected antigenic protein sequences were compared with representative human leukocyte antigen (HLA) class II alleles, available as a predefined set in IEDB (https://tools.iedb.org/mhcii/). Peptides with high binding affinity using percentile rank and predicted binding score were shortlisted. High affinity epitopes which potentially represent strong binders were prioritized to maximize the helper T cell activation.

### 2.3 Linear B-cell Epitope Prediction

Linear B-cell epitopes were predicted using the BepiPred 2.0 server available through the Immune Epitope Database (IEDB) (https://tools.iedb.org/bcell/), which employs a random forest machine-learning algorithm trained on experimentally validated epitope datasets to assign residue-level propensity scores. In contrast to previous propensity scale algorithms, BepiPred 2.0 combines sequence-based features with structural data to enhance the precision of the prediction of antibody-accessible regions[13]. This method allows the detection of peptide fragments that have a greater chance of being directly recognized by B-cell receptors.

Candidate B-cell epitopes were predicted based on their antigenicity, surface accessibility, and antibody recognition potential, which were predicted regions with high epitope probability scores. As B-cell epitopes are usually found in exposed and flexible parts of proteins, this filtering step ensures that the chosen peptides are structurally accessible to interact with immunoglobulins [13]. These epitopes were then taken into consideration to be included in the vaccine construct to induce humoral immune responses such as production of antibodies and neutralization of antigens.

### 2.4 Allergenicity Assessment

To achieve immunological safety, the predicted epitopes were assessed as allergenic potential using AllerTOP v.2 (https://www.ddg-pharmfac.net/AllerTOP/), an alignment-independent server, which uses auto- and cross-covariance transformation of amino acid properties and a machine-learning classifier. This approach determines physicochemical patterns linked to known allergens instead of using sequence similarity alone, which allows functional prediction of allergenicity. All epitopes were categorized as non-allergenic, which means that they are unlikely to cause IgE-mediated hypersensitivity reactions [14].

Filtering of allergenicity is critical in the design of multi-epitope vaccines because allergen-like peptides can trigger mast cell and basophil activation, which results in the release of histamine and other undesirable inflammatory responses. Only the epitopes that were predicted to be non-allergenic were selected to be analyzed downstream, so that the final construct would focus on immune activation without jeopardizing safety [14]. This, together with antigenicity and toxicity screening, aids in the selection of immunogenic and immunologically tolerable epitopes to be included in the proposed vaccine candidate.

### 2.5 Physicochemical properties, Toxigenicity and IL-10 response

The ProtParam web server (https://web.expasy.org/protparam/) was used to evaluate the physicochemical characteristics of the selected multi-epitope vaccine [15]. This program makes predictions on the molecular weight, theoretical isoelectric point (pI), instability index, aliphatic index, grand average of hydropathicity (GRAVY), extinction coefficient, and estimated protein half-life. The parameters assessed included molecular mass, protein stability profile, flexibility index, hydropathic nature and aliphatic index [16]. The theoretical charge behaviour was predicted via isoelectric point calculations. [16]. Furthermore, other sophisticated web-based tools were used to predict the vaccine’s toxigenicity and IL-10 response.

### 2.6 Multi-epitope vaccine construct

The multi-epitope vaccine construct was logically designed by arranging the selected B-cell, CTL and HTL epitopes corresponding to the three target proteins together in a single chimeric amino acid sequence. Epitope processing and presentation were improved by the use of AAY linkers for CTL epitopes to ensure efficient MHC-I epitope presentation and GPGPG linkers for HTL epitopes to preserve epitope structural flexibility and efficient MHC-II epitope presentation. Humoral immune recognition was also enhanced by introducing predicted linear B-cell epitopes in the construct. The overall epitope-linker arrangement was optimized for maximizing antigen processing efficiency, immunogenicity and coordinated activation of cellular and humoral immune responses using a proven PADRE adjuvant.

### 2.7 Primary and Secondary Structure Prediction

The full amino acid sequence of the assembled construct was firstly characterized using ExPASY ProtParam to determine its physiochemical properties and annotated to map all functional components across the full 823-residue chimeric sequence. The secondary structure of the vaccine construct was predicted using the PSIPRED Protein Sequence Analysis Workbench (http://bioinf.cs.ucl.ac.uk/psipred/), which achieves approximately 82 to 84% three-state (helix/strand/coil) secondary structure prediction accuracy [18]. PSIPRED classifies each residue into helix, strand, or coil, and also identifies disordered regions. The proportions of each structural class across the full 823-residue construct were recorded. The Alpha-helical and coil-rich regions are known to be associated with surface exposure and conformational flexibility, which support epitope accessibility and immune cell recognition [19].

### 2.8 Tertiary Structure Prediction, Refinement, and Validation

The three-dimensional(3D) structure of the vaccine construct was predicted at using the AlphaFold Server (https://alphafoldserver.com) at default parameters. This tool has shown great accuracy in predicting the complex configuration of proteins and other macromolecules [20]. This model works by apportioning per-residue confidence scores (pLDDT) and a predicted alignment error (PAE) matrix. The original predicted model was subjected to further refinement using 3DRefine. A disulfide bond engineering was performed using Disulfide by Design to increase the stability of the construct. A structural validation step was then conducted to validate the mutant model using PROCHECK. This program makes use of a Ramachandran plot and helps visualize the percentage of amino acid residues that are in favoured, allowed and disallowed regions. The next structural refinement involved the use of ProSA-web server (https://prosaservices.came.sbg.ac.at/prosa.php), which aids in calculating the overall quality Z-score of the mutant 3D model and comparing it to the score of protein structures obtained through X-ray analysis, NMR spectroscopy and theoretical calculations.

### 2.9 Molecular Docking of designed vaccine with TLR1 and TLR2

Immune response against mTB is strongly influenced by TLR1 and TLR2 mediated recognition of mycobacterial components, and TLR-based agonist have been widely explored as adjuvants to enhance the immunogenicity of modern TB vaccine formulation [21]. The TLR1/TLR2 heterodimer crystal structure (PDB: 2Z7X) was retrieved from the RCSB Protein Data Bank and prepared in PyMOL. The ClusPro v2.0 web server (https:cluspro.bu.edu/home.php) was used to docked with the disulfide-engineered vaccine construct as ligand and TLR1/TLR2 as receptors. Models were ranked by cluster size and weighted energy score and the highest ranked model for each docking prediction evaluated with PRODIGY (https://wenmr.science.uu.nl/prodigy/). PRODIGY helped to obtain the binding affinity, dissociation constant of our construct. iMODS web server (http://imods.chaconlab.org/) was used in the molecular dynamics assessing structural stability and conformational dynamics following molecular docking.

### 2.10 Construct Structural Validation

The stereochemical quality of the predicted three-dimensional (3D) structure of the vaccine construct was assessed using the PROCHECK server (https://www.ebi.ac.uk/thornton-srv/software/PROCHECK/). The Protein Data Bank (PDB) file of the model was uploaded to the server, and the structure was validated by examining the backbone dihedral angles (φ and ψ) in the Ramachandran plot. The number of residues in the most favoured, additionally allowed, generously allowed, and disallowed regions was noted [22]. Additional structural validations by PROCHECK included side-chain dihedral angles, bond lengths, bond angles, planar groups and residue properties. The stereochemical quality of the model was also evaluated using G-factors, which represent the normality of the covalent and torsional geometry of the structure. To assess the overall quality of the predicted structure, ProSA-web (https://prosa.services.came.sbg.ac.at/prosa.php) was employed[22]. The PDB file of the vaccine construct was submitted to the server, and the quality of the model was evaluated by the Z-score, which represent the difference between the total energy of the structure and the mean total energy of a database of experimentally determined protein structures. The Z-score obtained was compared with the Z-scores of native proteins of comparable size to assess the quality of the structure.

### 2.11 Normal Mode Analysis (NMA) of the Vaccine Construct

The dynamics and stability of the vaccine construct were predicted using the iMODS server (http://imods.iqf.csic.es), that uses Normal Mode Analysis (NMA) with an Elastic Network Model (ENM). The PDB file of the refined structure of the construct was uploaded for analysis. The server uses NMA based on internal coordinates (icNMA) to predict protein motions[23]. The deformability of residue was assessed to determine flexibility and hinge points, and the mobility profile was determined using calculated B-factors to estimate residue fluctuations. Eigenvalues for normal modes were calculated to predict stiffness, with smaller values suggesting less energy is needed for structural changes. Variance analysis was used to calculate the contribution of each mode to protein flexibility, including cumulative contributions. A covariance matrix was calculated to determine the dynamic interactions between residues, revealing correlated, uncorrelated and anti-correlated movements. Finally, we built an elastic network model to depict residues as spring-connected particles, to assess structural network connectivity and system stiffness.

### 2.12 Disulfide Engineering of the Vaccine Construct

Disulfide bond engineering of the predicted vaccine construct was carried out using Disulfide by Design 2 server from Wayne state university (http://cptweb.cpt.wayne.edu/DbD2/). The server was provided with the 3D structure of the vaccine construct in PDB format. Residue pairs that could form disulfide bonds were identified according to geometric parameters such as the distance between the Cβ atoms and the χ3 dihedral angle. Residue pairs meeting the conventional criteria for disulfide bond formation were mutated to cysteine residues. The mutant structures were examined and visualized to ensure the formation of disulfide bonds and to check the feasibility of the structures[24].

### 2.13 In Silico Immune Simulation

The immunogenicity of the constructed multi-epitope vaccine construct was assessed with the help of the C-IMMSIM server (http://kraken.iac.rm.cnr.it/C-IMMSIM/), a computational platform based on metrics simulating mammalian immune responses through the modelling of interactions between immune cells, cytokines, and antigen presentation pathways. The simulation includes important immunological events, such as antigen uptake, MHC class I and II presentation, lymphocyte clonal expansion, and cytokine-mediated signaling. The model behind it is position-specific scoring matrices and immune system dynamics as outlined by [25]. The default parameters were used to perform simulations, and various antigen exposures at an approximate a thousand molecules and 10µl were given at specific time between day 1 prime and day 60 boost, following steps to simulate prime boost vaccination. Humoral and cellular immune responses such as immunoglobulin production (IgM, IgG subclasses), B-cell population dynamics, CD4+ and CD8+ T cell responses, cytokine profiles (especially IFN-g and IL-2), and memory cell formation were assessed in the model [26]. The immune response curves were examined to determine primary, secondary and tertiary responses, antigen clearance and long-term immunological memory, which gave a complete in silico validation of the immunogenicity of the vaccine construct.

## 3.0 RESULTS

### 3.1 Linear B-cell Epitope Prediction

Hydrophilicity-based prediction techniques were used to identify linear B-cell epitopes, and immunological screening was performed. Several candidate epitopes were predicted, and the top-ranking peptides were chosen on the basis of antigenicity, safety and immunomodulatory potential (Table 1). The epitopes that were chosen had high scores of antigenicity with a maximum score of 1.6061, which means that they are highly immunologically recognized. The shortlisted epitopes were non-allergenic and non-toxic, which validated their use in the development of multiple-epitope vaccines. Notably, most of the identified B-cell epitopes had a positive interferon (IFN) induction potential, indicating their ability to induce Th1-biased immune responses, which are essential in the defense against intracellular pathogens like Mycobacterium tuberculosis. Also, the lack of IL-10 induction implies a decreased risk of anti-inflammatory suppression, which also contributes to successful immune activation. Representative epitopes across the three proteins were sequences like RPNGQAGSN and RYGIDNEPQGVAGGGKAVEALGLNPPPVP that were high in antigenicity scores and had good immunological characteristics. These results indicate the possibility of the chosen B-cell epitopes to play a role in the development of strong humoral immunity.

**Table 1.**
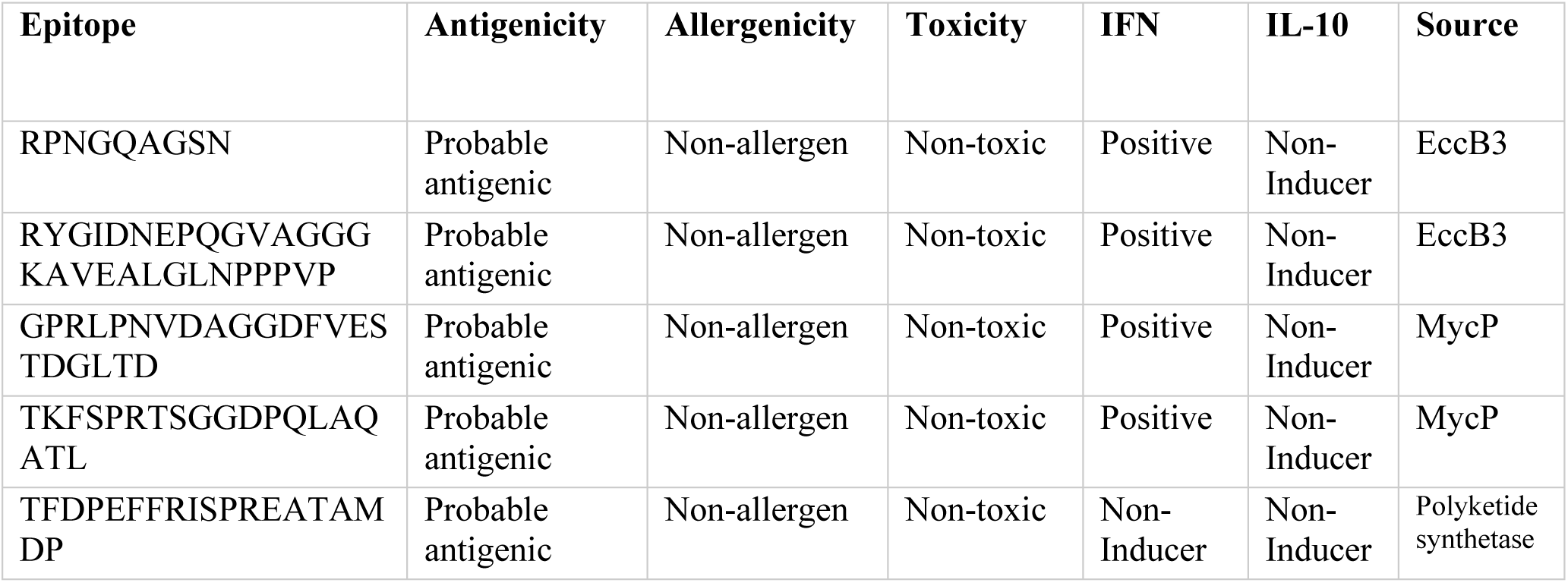
Selected B-cell epitopes and their immunological properties.

### 3.2 Cytotoxic T Lymphocyte (CTL) Epitope Prediction

The prediction of cytotoxic T lymphocyte (CTL) epitopes was done to determine the peptide sequences that could bind MHC class I molecules and trigger CD8+ T cell-mediated immune responses. A complete repertoire of candidate epitopes was produced and then narrowed down on the basis of antigenicity, safety and immunogenicity (Table 2). The chosen CTL epitopes were shown to have high binding affinity in various HLA class I alleles, which means that they have a high potential to be presented to CD8+ T cells. It is worthy to note that all selected epitopes had good antigenicity scores, which implies that they can be used to induce immune recognition. All the shortlisted epitopes were predicted to be non-allergenic and non-toxic, which means that they would be safe for a vaccine inclusion.

**Table 2.**
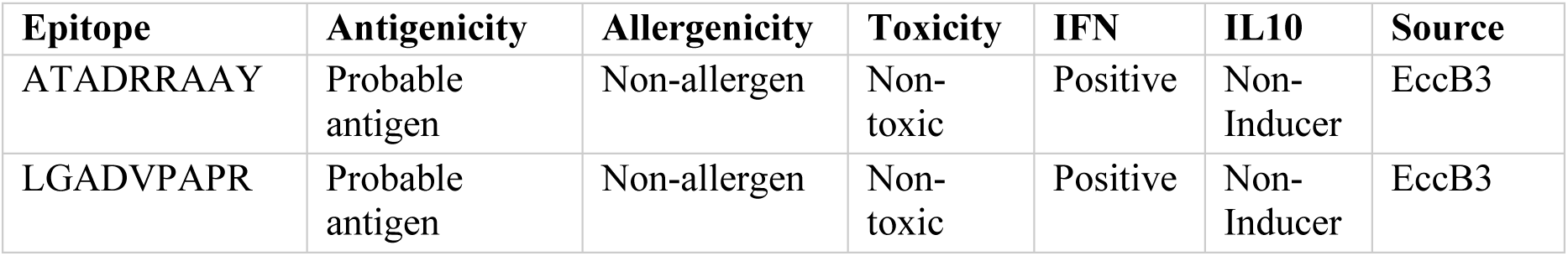
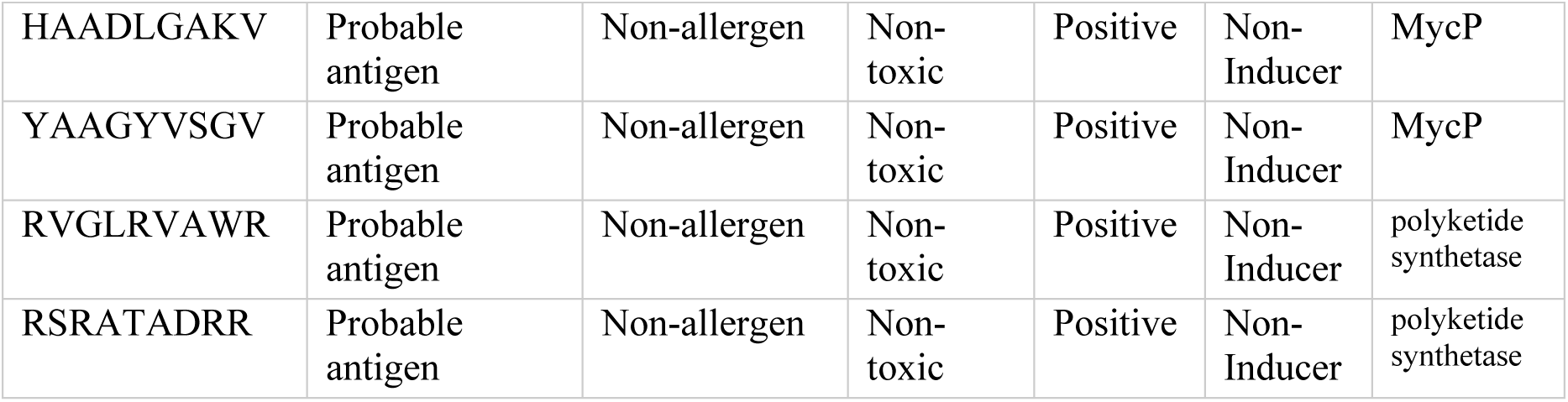
presents a detailed list of the selected CTL epitopes, their respective sources and immunological properties.

Moreover, the diversity of epitopes in various HLA alleles underscores their possible wide population coverage, which enhances the chances of immune responsiveness in various genetic backgrounds. These CTL epitopes are essential in the induction of cell-mediated immunity which is central in the elimination of intracellular pathogens like Mycobacterium tuberculosis by facilitating the recognition and destruction of infected host cells. The representative high-affinity epitopes that were found in the current study were peptides like ATADRRAAY, LGADVPAPR, and LHVDEPSRE that exhibited good binding affinities with various HLA alleles and good immunogenicity.

### 3.3 Helper T Lymphocyte (HTL) prediction

The prediction of helper T lymphocyte (HTL) epitopes was done to determine the peptide sequences that could bind the MHC class II molecules and trigger the immune response mediated by CD4+ T cells. An extensive list of candidate epitopes was produced and then narrowed down according to antigenicity, allergenicity, toxicity and cytokine induction potential (Table 3). The chosen HTL epitopes were found to be of high binding affinity to various HLA-DR alleles, such as DRB10101 and DRB10301, which suggests that they may be widely recognized by the immune system of different populations. Selected epitopes had high antigenicity scores, which validated their appropriateness in the generation of strong immune responses. Notably, shortlisted HTL epitopes were all predicted to be non-toxic, non-allergenic, and demonstrated positive interferon (IFN-γ) induction capacity. The representative HTL epitopes were LHVDEPSRE, FRISPREAT and LGADVPAPR, which showed good binding affinity and good immunogenicity.

**Table 3.** gives a detailed list of the chosen HTL epitopes, their immunological characteristics, and their sources.

| Epitope | Antigenicity | Allergenicity | Toxicity | IFN-γ | IL 10 | SOURCE |
| --- | --- | --- | --- | --- | --- | --- |
| LHVDEPSRE | Probable antigen | Non-allergen | Non-toxic | Positive | Non-Inducer | EccB3 |
| FRISPREAT | Probable antigen | Non-allergen | Non-toxic | Positive | Non-Inducer | EccB3 |
| IRAAFACLAATVVVA | Probable antigen | Probable Non-allergen | Non-Toxic | Positive | Non-Inducer | MycP |
| LAGAIVHAADLGAKV | Probable antigen | Probable non-allergen | Non-Toxic | Positive | Non-Inducer | MycP |
| VGCYVGASALEYGPA | Probable antigen | Probable non-allergen | Non-Toxic | Positive | Non-Inducer | polyketide synthetase |
| DVGCYVGASALEYGP | Probable antigen | Probable non-allergen | Non-Toxic | Positive | Non-Inducer | polyketide synthetase |

### 3.4 Multi-epitope vaccine construct design

The final multi-epitope vaccine construct was assembled by integrating selected B-cell, CTL, and HTL epitopes from the three (3) proteins into a single chimeric sequence. This construct featured 11 MHC I, 14 MHC II and 17 B cell epitopes. The CTL epitopes were linked using AAY and KK linkers while HTL epitopes alongside B cell epitopes were joined using GPGPG linkers. These glycine abundant linkers were employed due to their profound solubility and flexibility to preserve epitope integrity and enhance MHC class II presentation. Furthermore, the N and C -terminal of the multi-epitope construct were joined with the EAAAK and His-tag linkers respectively (Figure 2A). This rational arrangement of epitopes and linkers was designed to ensure optimal antigen processing, presentation, and immune activation. The final construct sequence is contained in the supplementary data.

**Figure 2.**
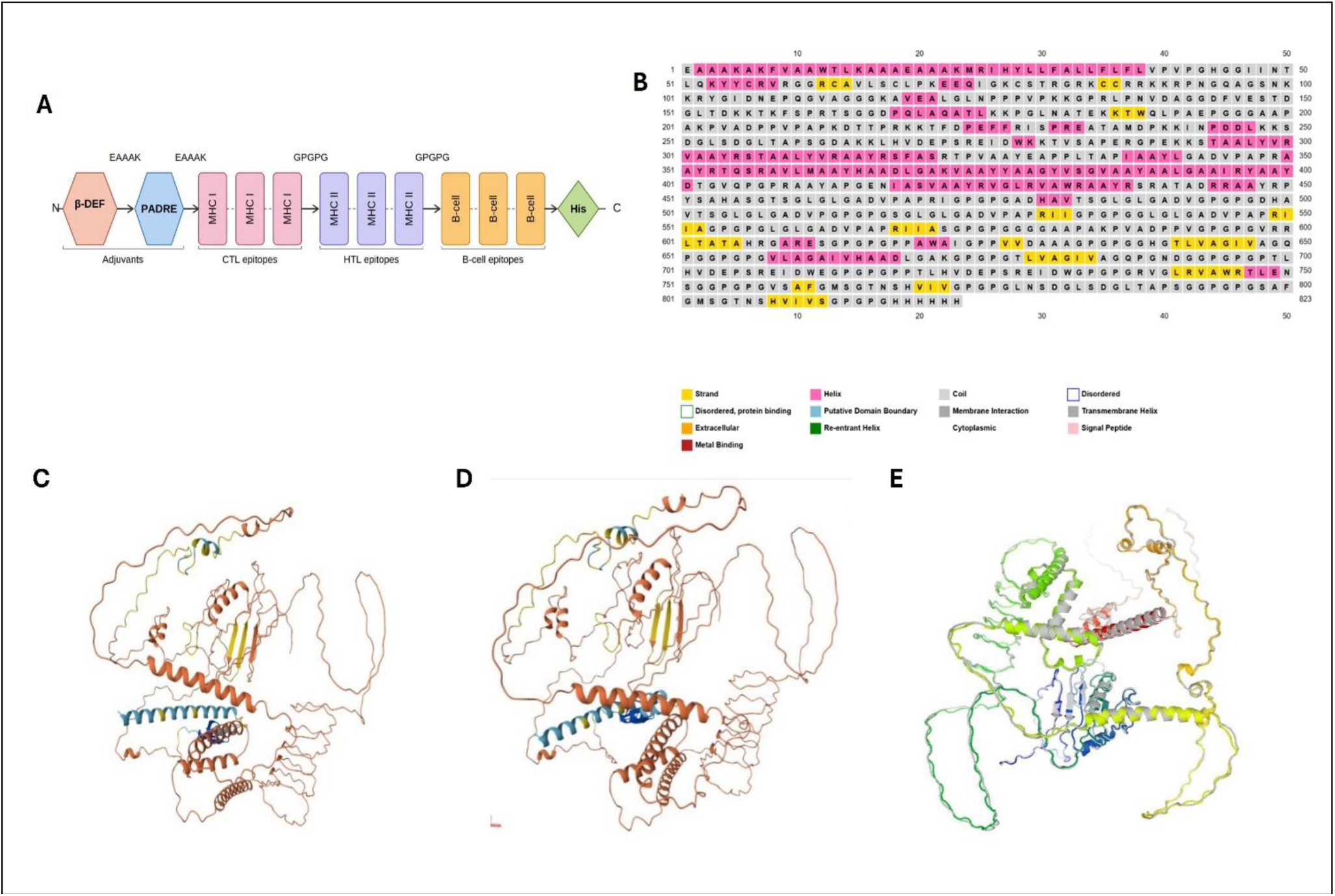
Sequence architecture and predicted structure of the multi-epitope vaccine construct. (A) The linear architecture shows the arrangement of all functional components from N- to C-terminus. (B) PSIPRED secondary structure prediction output for the 823-residue multi-epitope vaccine construct. **(C**–**D)**: Predicted tertiary structure showing the alpha-helical core and flexible linker coils. **(E)**: Refined tertiary structure following 3DRefine processing with improved side-chain geometry and reduced local steric strain.

### 3.5 Physicochemical characteristics

The multiple epitope vaccine construct consists of 822 amino acids, with a molecular weight of approximately 82.3kDa. The theoretical PI was 9.61 which indicates that it is strongly basic, with an instability index of 32.48 indicating the protein is stable under normal physiological conditions. The Gravy of -0.263 predicts that the multi-epitope vaccine has overall hydrophilicity, therefore it is very soluble. Furthermore, it has an aliphatic index of 72.48 which reflects moderate thermostability. This vaccine construct’s half-life was predicted to be 1-hour in vitro mammalian reticulocyte, over 30 minutes in yeast and more than 10 hours in *E. coli* indicating the protein is potentially stable in diverse expression systems.

### 3.6 Secondary Structure Analysis

PSIPRED analysis of the 823-residue construct predicted a secondary structure composition of 20.5% alpha-helix (169 residues), 5.3% beta-strand (44 residues), 49.8% coil (410 residues), and 24.3% disordered regions (200 residues), as shown in Figure 2B. The high proportion of coil and disordered residues (74.1% combined) is consistent with the architecture of the construct, which incorporates multiple GPGPG and AAY flexible linker sequences that are deliberately disordered to preserve conformational independence of adjacent epitopes during antigen processing and presentation. The alpha-helical content (20.5%) reflects the structured beta-defensin adjuvant at the N-terminus, which is known to adopt a helical conformation, alongside helical regions within the CTL and HTL epitopes. The low beta-strand content (5.3%) of the construct is broadly consistent with published multi-epitope TB vaccine designs, which often show secondary structures dominated by random coil with relatively limited β-strand content. [27].

### 3.7 Tertiary Structure Prediction, Refinement, and Validation

AlphaFold prediction produced a model with a mixed alpha-helical and coil topology with localized beta-strand content (Figure 2C**–**D), consistent with the PSIPRED secondary structure prediction. Structured core regions showed confident pLDDT scores (70-90), while linker regions displayed lower confidence (below 70), reflecting their designed intrinsic flexibility. The pTM score of 0.17 is expected for an artificial chimeric sequence with no natural structural homologue in AlphaFold’s training data, and model quality is assessed independently through the validation metrics below. The model was refined using 3DRefine to improve hydrogen bonding geometry and side-chain conformations as shown in Figure 2E.

### 3.8 Disulfide bond engineering

The Structural comparison of the native and mutant models showed the formation of disulfide bonds at the desired residue positions. The mutant model showed the formation of covalent bonds between the cysteine residues, as shown in the mutant structure (Figure 3A), suggesting that the chosen pairs of residues met the geometric and conformational constraints for disulfide bond formation. These include favourable χ3 dihedral angles and inter-residue distances, allowing the formation of the disulfide bonds in the protein structure. The addition of these disulfide bonds is likely to increase the rigidity and stability of the vaccine construct by restricting its conformational flexibility. This is crucial for multi-epitope vaccines, which may include flexible linkers that can affect the structure. In summary, the disulfide engineering strategy led to a structurally refined and stabilized model, serving an assurance on improved stability, folding and potential immunogenic presentation of our vaccine construct.

**Figure 3.**
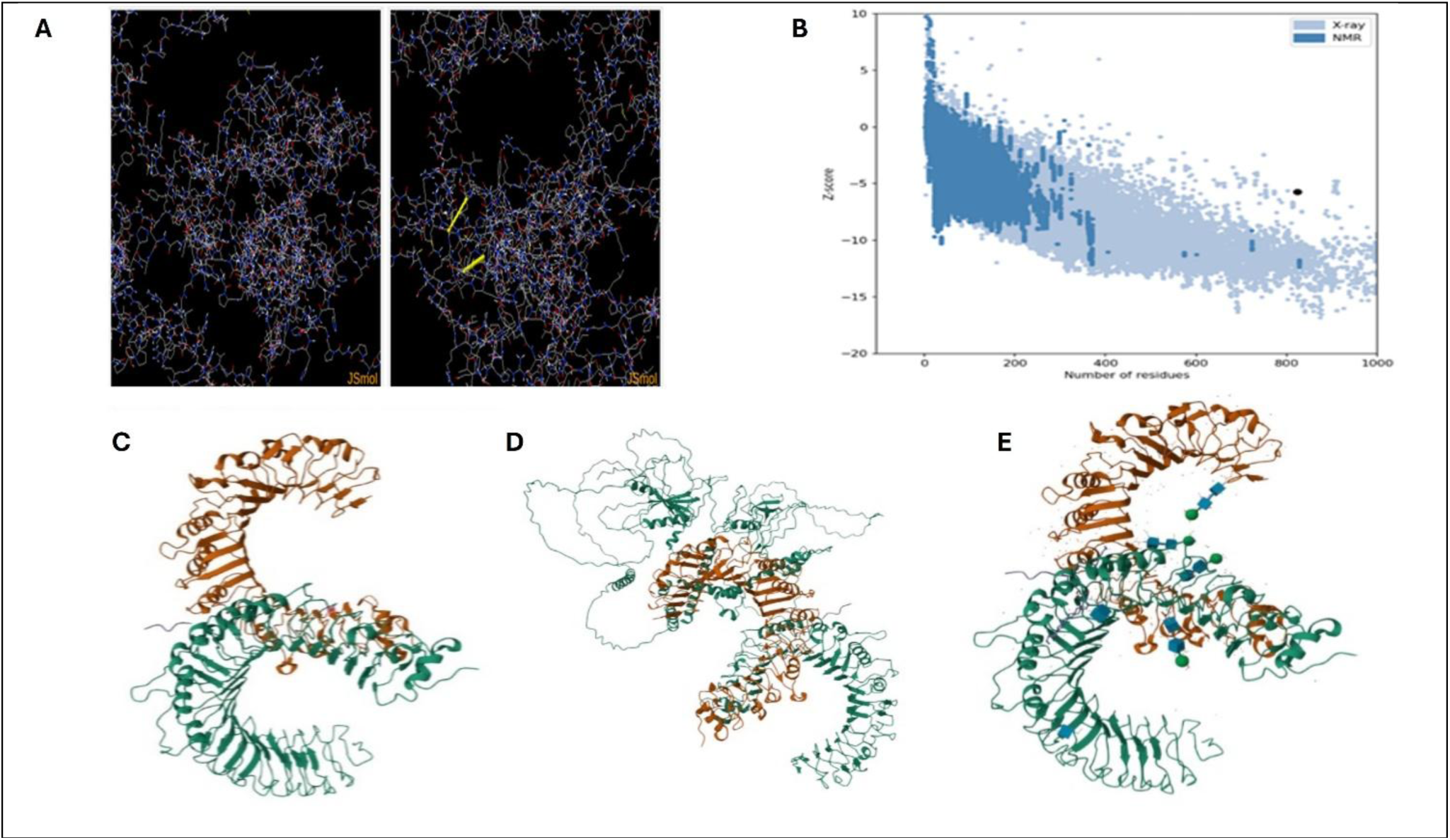
Disulfide engineering, structural validation, and TLR1/TLR2 docking of the vaccine construct. **(A)** The disulfide bond (in yellow) as predicted by Disulfide by Design 2, **(B)** The Z-score (-6) obtained with ProSa-Web. (C) TLR1/TLR2 heterodimer receptor structure used as the docking target before ligand removal. **(D)**: Docked complex of the vaccine construct (orange) and TLR1/TLR2 heterodimer (green) after ligand removal. **(E)**: Binding pocket bound to the vaccine construct.

### 3.9 Structural validation

The stereochemical properties of the predicted three-dimensional model were assessed using the PROCHECK server. The Ramachandran plot showed 94.7% of residues in the most favored regions, 4.1% in additionally allowed regions, 0.5% in generously allowed regions and 0.7% in disallowed regions. This suggests that most backbone dihedral angles are in energetically preferred conformations, validating the model. Also, the model displayed overall bond lengths, bond angles and planar groups, which were all within the expected range, with 100% of planar groups within the expected range. The overall G-factor (∼0.14-0.15) also confirmed the quality of the structure, with values above -0.5 being considered good stereochemistry.

Further validation of the overall quality of the model using the ProSA-web server presented a Z-score of −6, comparable to those observed for naturally occurring and experimentally validated proteins of comparable size via X-ray and NMR (Figure 3B). These findings show that the predicted tertiary structure of the vaccine construct is stable and of good quality, and much suited for further downstream evaluation.

### 3.10 Molecular Docking with the TLR1/TLR2 Heterodimer

Molecular docking evaluates the interaction between the ligand molecule (the refined mutant vaccine structure) and Toll-like receptor molecules (TLRs 1 and 2) in their optimal complex conformations. This technique reveals the binding affinity between the molecules through a scoring function and provides valuable insights into their interaction. ClusPro 2.0 generated 333 docked models across 23 clusters. The top-ranked model (Cluster 0) selected had the largest cluster size of 32 members and a weighted energy score of -1371.0 kcal/mol. The docked complex reveals a well-defined interface between the vaccine construct and the TLR1/TLR2 ectodomain, with extensive contacts along the concave leucine-rich repeat domain surface (Figure 3D–E).

### 3.11 Normal mode analysis and structural dynamics

Normal mode analysis revealed the vaccine construct has a stable structure with flexible regions. The deformability profile exhibited low variability for most residues, suggesting a relatively rigid structure, but also displayed significant peaks, especially around residues ∼1000-1400, which suggest the presence of deformable regions that can act as hinges to enable conformational transitions (Figure 4A). The eigenvalue for the first normal mode (2.27 × 10⁻⁸) indicated that the structure is easily deformable, which is indicative of a flexible structure with a good stability-flexibility balance. Variance analysis showed that most of the motion is explained by the first few normal modes, with variance quickly reaching a plateau (Figure 4D). This suggests that protein motions are dominated by collectives rather than random motion. The covariance matrix displayed regions of positive and negative covariance between residues, suggesting intra-molecular communication and domain coordination (Figure 4F). Additionally, the elastic network model revealed a highly connected network, providing structural integrity and allowing local deformations (Figure 4E). Overall, these findings demonstrate that the construct has favourable dynamic characteristics, achieving the structural stability and flexibility needed for biological function.

**Figure 4.**
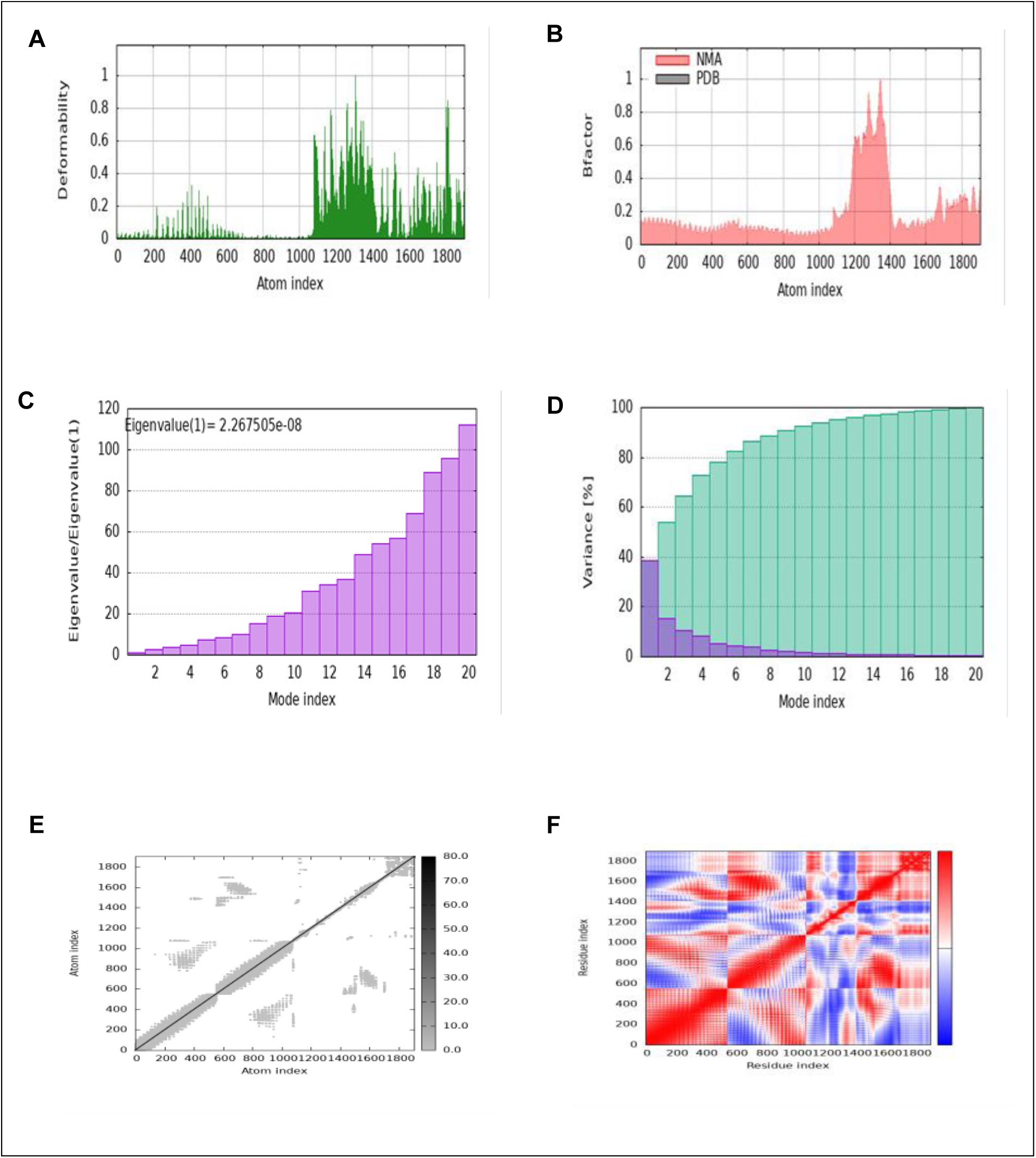
Normal mode (structural dynamics) analysis of the vaccine–TLR1/TLR2 docked complex. (**A**) Deformability plot showing the flexibility of various regions within the docked complex (**B**) B-factor analysis representing atomic fluctuations across the complex. (**C**) Eigenvalue analysis depicting the stiffness of the docked structure and its associated energy requirements. (**D**) Variance analysis representing the structural variations in the docked complex. (**E**) Elastic network analysis illustrates the connectivity and motion of residues within the complex (**F**) Covariance analysis highlighting the correlated motions of residues.

### 3.12 In Silico immune simulation

The C-IMMSIM platform was used to assess the immunogenicity of the multi-epitope vaccine construct after simulated primary and booster vaccinations. The model showed successful antigen uptake and presentation by antigen-presenting cells (APCs) after the initial dose, which resulted in the activation of both CD4+ and CD8+ T lymphocytes. There was a distinct peak of IL-2 production, which is in line with T-cell proliferation and early immune activation. The humoral response was typical primary pattern, with an initial increase of IgM antibodies, and a gradual increase of IgG. At the same time, B-cell populations increased and started to shift towards memory states, which is a sign of successful immune priming without any signs of overreacting inflammatory reactions. Following the second (booster) dose, a significantly stronger immune response was detected, indicating successful immunological memory formed in the course of the priming stage (Figure 5).

**Figure 5.**
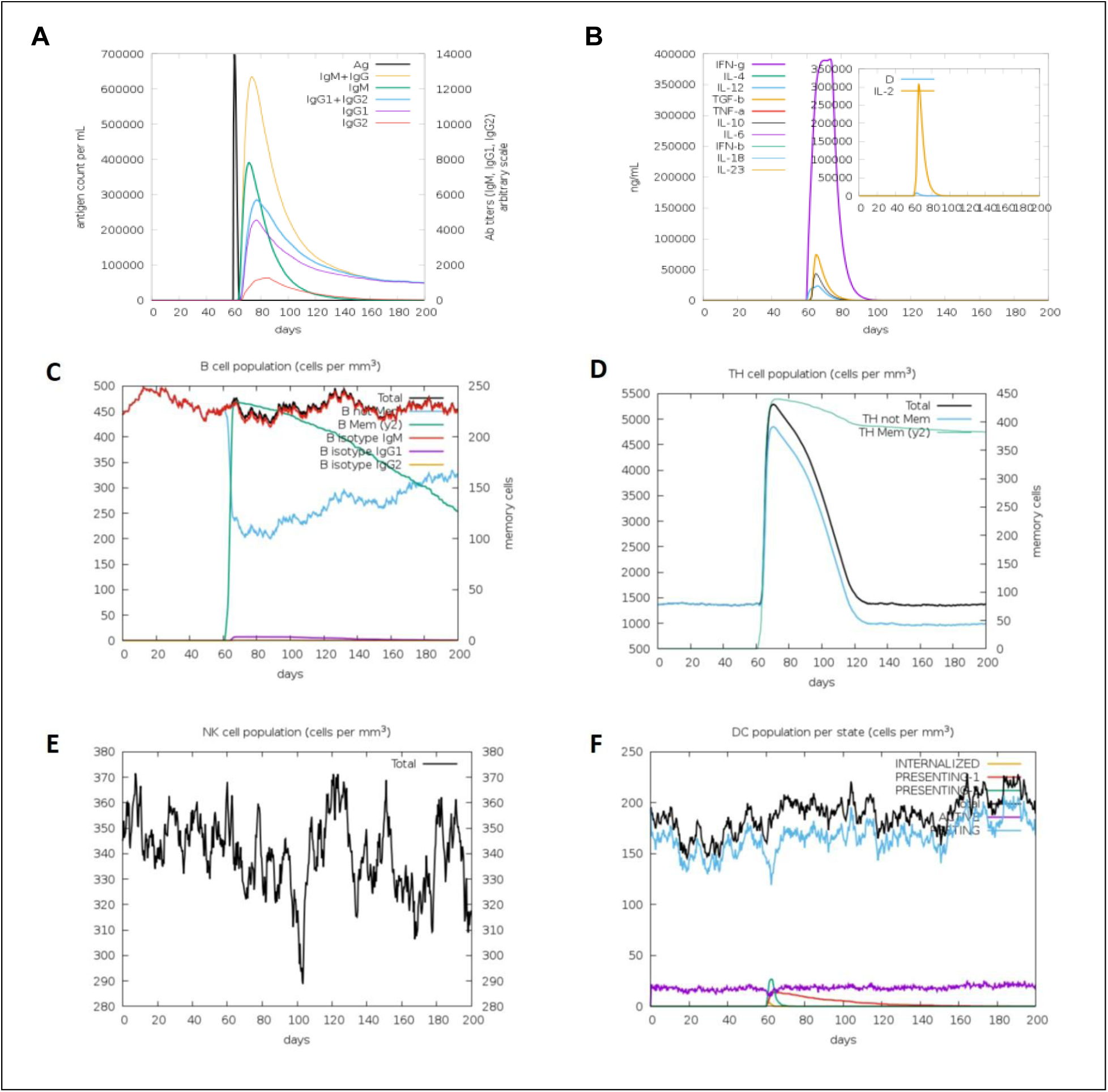
C-ImmSim simulation of the immune response to the vaccine construct. **(A)** The test virus, the immunoglobulins and the immunocomplexes after the second dose, **(B)** Concentration of cytokines and interleukins. The Inset plot shows no danger signal together with leukocyte growth factor IL-2 following the booster dose. **(C)** B cell population showed assertive increment across all subunits, **(D)** Helper T cells responding to dose with high titer value over period of few months before dropping down to initial levels, **(E)** NK cells population showed somewhat erratic but generally good response to vaccine, **(F)** Dendritic cells population maintained uniform peak throughout signifying proper innate immune response.

The antibody kinetics revealed a rapid and strong rise in the level of IgG, which is much higher than the primary response, classical evidence of class switching and affinity maturation. B-cell dynamics showed a steep increase in active B cells and then a stabilization into a long-lasting pool of memory B cells. Likewise, the CD4+ T-helper cells were highly activated with stable resting populations whereas the CD8+ cytotoxic T cells grew and then shifted into a stable memory pool. This synchronized reaction attests to effective recall immunity and increased responsiveness to re-exposure to the antigen (Figure 5). The cytokine profiling of both doses showed a Th1-biased immune response with significant peaks of IFN-g and IL-2 after antigen exposure, which promotes T-cell proliferation and intracellular pathogen control. Similarly, the profiles of NK and DCs returned optimum for a good innate immune sensing. The booster dose resulted in a faster clearance of the antigens, which also suggested better immune efficiency. The existence of long-term memory B- and T-cells, along with strong humoral and cellular immunity, indicates that the vaccine construct can induce long-term protective immunity. In general, the simulation demonstrates an effective shift in the primary immune activation to a robust secondary response, which confirms the immunogenicity of the developed multi-epitope vaccine candidate.

## 4.0 DISCUSSION

This study has adopted a comprehensive immunoinformatics approach to design a multi-epitope vaccine against major virulence factor proteins of Mycobacterium tuberculosis (Mtb) - EccB3, MycP and polyketide synthase[28]. These were selected for their key roles in nutrient acquisition, secretion system function and immune resistance. This design strategy is in line with current research on TB vaccines, where multi-antigen targeting has been shown to have better immunogenicity and lower evasion ability. The unique feature of this study is the selection of highly antigenic, immune-evasive and non-toxic epitopes capable of inducing humoral and cellular immune responses [29]. The predicted B-cell epitopes had high antigenicity and the targeted immunogenicity, including positive induction of IFN and negative induction of IL-10, which predicts a Th1-bias immune response [30]. This is essential in the induction of immunity to TB, which relies on activation of an immune response that results in the activation of macrophages to kill intracellular pathogens through IFN-γ [30]. Similarly, the CTL and HTL epitopes demonstrated high affinity to a wide range of HLA alleles suggesting that it will be able to reach a large number of people and induce antigen presentation via both MHC class I and II [29]. These findings are consistent with recent research that emphasizes the use of multi-epitope constructs in driving both CD4⁺ and CD8⁺ T cell responses against intracellular pathogens [30].

The structure of the vaccine construct is also in tandem with its potential use as a vaccine. The construct was found to be stable, and soluble. The high proportion of coil and disordered structure in the secondary structure prediction is consistent with the design of flexible linkers (AAY and GPGPG) to enhance the processing and presentation of the epitopes. This helps in the recognition of antigens by B-cell receptors, and presentation by antigen-presenting cells via MHC molecules. The tertiary structure prediction by AlphaFold, refinement and validation have shown that the construct has a good chimeric structure. Structural validation, examined in the Ramachandran plot and ProSA Z-scores, revealed that the construct is similar to those with experimentally determined structures, and likely has a good stereochemistry [31]. Minor differences in side-chain positions, as seen in large synthetic proteins, do not affect the structure.

Molecular docking simulations revealed strong and stable interaction of the vaccine construct with the TLR1/TLR2 heterodimer, with a deep and highly negative docking energy and a clear docking interface. This interaction is important as TLR1/TLR2 is recognized to be essential for the recognition of mycobacterial antigens and subsequent activation of the innate immune response [32]. This engagement suggests an intrinsic adjuvant effect of the vaccine construct, which in turn could result in enhanced antigen presentation and, thus, adaptive immunity. This finding is consistent with recent reports that vaccine constructs that can directly engage pattern recognition receptors, thereby increase immunogenicity [15].

Moreover, the flexibility of the vaccine construct, as predicted by normal mode analysis, highlights its potential. The Eigenvalue is low, implying flexibility, while the rigid and deformable regions demonstrate stability and flexibility. These are essential for binding and immunogenicity. The covariance and elastic network analyses also revealed correlated motions of residues, indicating the stability and functionality of the product [20].

The design of disulfides provided additional stability through covalent bonding of some residues. This is especially important in multi-epitope vaccines that have flexible regions that can reduce rigid folding[33]. The capacity of the mutant model to form disulfide bonds suggests greater stability and folding, which is crucial for vaccine stability and effectiveness. More importantly, the in-silico immune simulation demonstrated the immunogenicity of the vaccine. It demonstrated the typical initial immune response with an IgM response, and a robust secondary response with a significant increase in IgG following the booster dose[34]. This suggests class switching and maturation, hallmarks of humoral response. Further, the presence of memory B and T cells indicate the establishment of immunological memory[26]. The high levels of Th1 cytokines (IL-2 and IFN- γ) are particularly encouraging, as they are linked to protective immunity against Mtb. Further, the rapid removal of antigen after the booster dose suggests efficient and memory responses of the immune system.

These findings align with previous studies, which indicate multi-epitope vaccines can revitalize natural immune responses and offer long-lasting immunity [12]. The ability of the C-IMMSIM simulation software to model key immune processes like antigen processing, cell activation and cytokine crosstalk also indicate the reliability of the predictions [26]. While these results are promising, there are some caveats. As an in-silico study, the findings are predictions which need to be verified through experimentation [6]. Also, while the construct has high binding affinity to TLR1/TLR2, it needs to be experimentally evaluated for its potential to trigger innate immune responses.

## 5.0 CONCLUSION

In this study, we established a rational design of a novel multi-epitope vaccine construct against major virulence proteins of Mycobacterium tuberculosis (Mtb) - EccB3, MycP and polyketide synthase - using immunoinformatics. The multi-epitope construct addresses multiple virulence factors, providing a more effective approach in comparison to the traditional single-antigen vaccines. The B-cell, CTL, and HTL epitopes were found to be highly antigenic, had good HLA coverage and were safe, suggesting the potential to elicit both humoral and cellular immunity. Structural and physicochemical studies validated the stability, solubility and flexibility of the construct, facilitating antigen processing. Refinement and validation of the tertiary structure also confirmed the model’s validity. Docking studies showed binding to the TLR1/TLR2 heterodimer, implying possible interaction with the innate immune system. Normal mode analysis showed good stability-flexibility, with disulfide engineering further increasing stability. Immune simulations demonstrated high immunogenicity, with efficient antigen presentation, robust secondary responses and durable memory responses. The Th1-dominated cytokine profile (IFN-γ and IL-2) is essential to combat intracellular pathogens, while IgG responses are enhanced on repeated exposure to the booster. In summary, the vaccine construct has good potential as a prophylactic solution for tuberculosis. However, experimental testing will be needed to assess its efficacy and safety in vivo.

---

**Supplementary Data 1:**
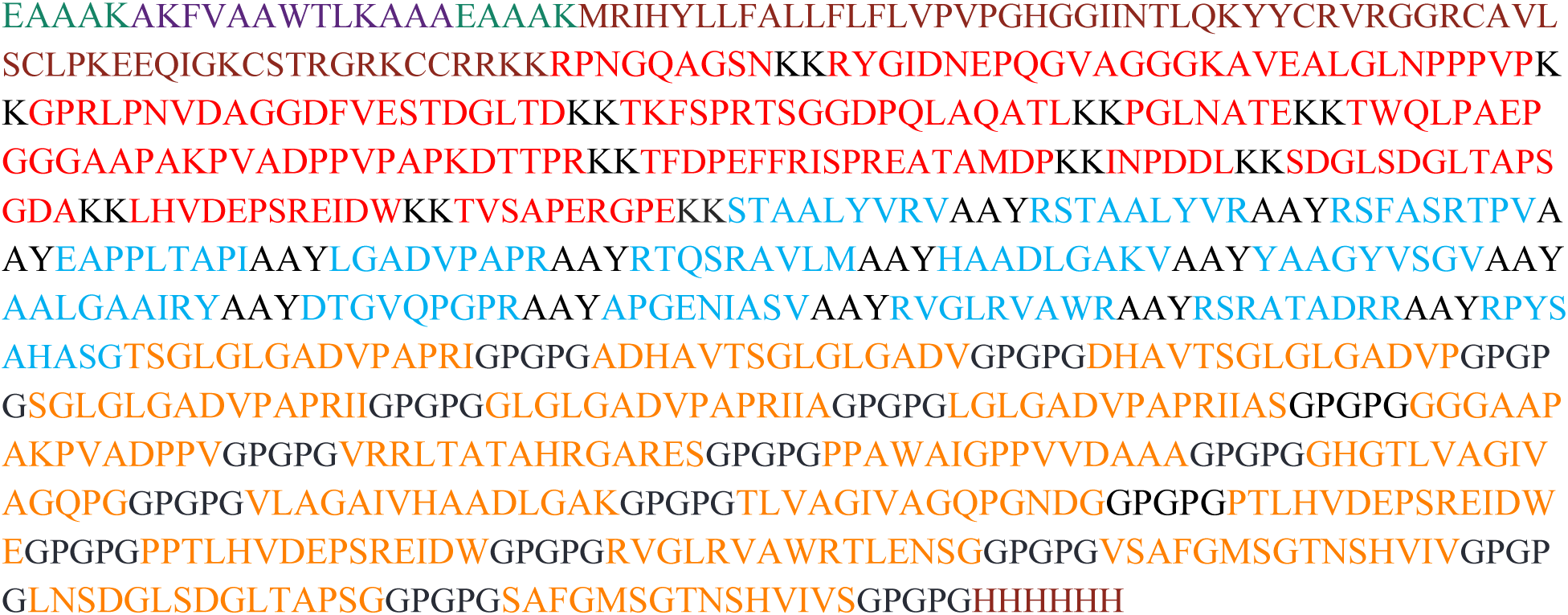
*Sequence showcasing the assembly of the amino acids chain of the vaccine construct*.

## REFERENCE

[1] Global Tuberculosis Report 2023. World Health Organization; 2023.

[2] Mancuso G, Midiri A, De Gaetano S, Ponzo E, Biondo C. Tackling Drug-Resistant Tuberculosis: New Challenges from the Old Pathogen Mycobacterium tuberculosis. Microorganisms 2023;11. 10.3390/microorganisms11092277.

[3] Passos BBS, Araújo-Pereira M, Vinhaes CL, Amaral EP, Andrade BB. The role of ESAT- 6 in tuberculosis immunopathology. Front Immunol 2024;15. 10.3389/fimmu.2024.1383098.

[4] Yang Y, Chen J, Liu L, Li L, Yang R, Lu K, et al. Applying a Combined Model to Evaluate the Risk of Poor Treatment Outcomes in Rifampicin Resistant Tuberculosis Patients: A Multicenter Retrospective Study. Infect Drug Resist 2024;17:5287–98. 10.2147/IDR.S491910.

[5] Sethi G, Varghese RP, Lakra AK, Nayak SS, Krishna R, Hwang JH. Immunoinformatics and structural aided approach to develop multi-epitope based subunit vaccine against Mycobacterium tuberculosis. Sci Rep 2024;14. 10.1038/s41598-024-66858-5.

[6] He Y, Rappuoli R, De Groot AS, Chen RT. Emerging vaccine informatics. J Biomed Biotechnol 2010;2010. 10.1155/2010/218590.

[7] Kawai T, Ikegawa M, Ori D, Akira S. Decoding Toll-like receptors: Recent insights and perspectives in innate immunity. Immunity 2024;57:649–73. 10.1016/j.immuni.2024.03.004.

[8] Bythrow G V., Farhat MF, Levendosky K, Mohandas P, Germain GA, Yoo B, et al. Mycobacterium abscessus Mutants with a Compromised Functional Link between the Type VII ESX-3 System and an Iron Uptake Mechanism Reliant on an Unusual Mycobactin Siderophore. Pathogens 2022;11. 10.3390/pathogens11090953.

[9] Bunduc CM, Fahrenkamp D, Wald J, Ummels R, Bitter W, Houben ENG, et al. Structure and dynamics of a mycobacterial type VII secretion system. Nature 2021;593:445–8. 10.1038/s41586-021-03517-z.

[10] Cambier CJ, Falkow S, Ramakrishnan L. Host evasion and exploitation schemes of Mycobacterium tuberculosis. Cell 2014;159:1497–509. 10.1016/j.cell.2014.11.024.

[11] Oladipo EK, Ajayi AF, Onile OS, Ariyo OE, Jimah EM, Ezediuno LO, et al. Designing a conserved peptide-based subunit vaccine against SARS-CoV-2 using immunoinformatics approach. In Silico Pharmacol 2021;9. 10.1007/s40203-020-00062-x.

[12] Olatunde SK, Oyedele AO, Alli YA, Oladipo EK, Ajisafe HM, Abiodun CS. Designing a multi-epitope vaccine to harness computational immunology for next-generation tuberculosis control. Next Research 2026;6:101398. 10.1016/j.nexres.2026.101398.

[13] Akurut E, Gavamukulya Y, Mulindwa J, Isiagi M, Galiwango R, Bbuye M, et al. Design of a multi-epitope vaccine against drug-resistant mycobacterium tuberculosis and mycobacterium bovis using reverse vaccinology. Sci Rep 2025;15. 10.1038/s41598-025-11768-3.

[14] Oladipo EK, Adeyemo SF, Oshoneye AI, Akintola HB, Elegbede BI, Ayoomoba TU, et al. Harnessing computational immunology to design targeted subunit vaccines for infectious bursal disease in poultry. Frontiers in Bioinformatics 2025;5. 10.3389/fbinf.2025.1562997.

[15] Francisco S, Billod JM, Merino J, Punzón C, Gallego A, Arranz A, et al. Induction of TLR4/TLR2 Interaction and Heterodimer Formation by Low Endotoxic Atypical LPS. Front Immunol 2022;12. 10.3389/fimmu.2021.748303.

[16] Naveed M, Asim M, Aziz T, Athar A, Majeed MN, Tombozara N, et al. In silico design and immunoinformatics assessment of a multiepitope vaccine targeting borealpox virus. Sci Rep 2026. 10.1038/s41598-026-36680-2.

[17] The Proteomics Protocols Handbook. n.d.

[18] Buchan DWA, Jones DT. The PSIPRED Protein Analysis Workbench: 20 years on. Nucleic Acids Res 2019;47: W402–7. 10.1093/nar/gkz297.

[19] Jia Y, Yang K, Sun Q, Guo W, Yang Z, Duan Z, et al. Design and Evaluation of a Broadly Multivalent Adhesins-Based Multi-Epitope Fusion Antigen Vaccine Against Enterotoxigenic Escherichia coli Infection. Vaccines (Basel) 2025;13. 10.3390/vaccines13101057.

[20] Abramson J, Adler J, Dunger J, et al. Accurate structure prediction of biomolecular interactions with AlphaFold 3. Nature. 2024;629(8012):493–500. doi: 10.1038/s41586-024-07487-w

[21] Junqueira-Kipnis, A. P., Marques Neto, L. M., & Kipnis, A. (2014). Role of fused *Mycobacterium tuberculosis* immunogens and adjuvants in modern tuberculosis vaccines. Frontiers in Immunology, 5, 188. 10.3389/fimmu.2014.00188

[22] Evangelista FMD, van Vliet AHM, Lawton SP, Betson M. In silico design of a polypeptide as a vaccine candidate against ascariasis. Sci Rep 2023;13. 10.1038/s41598-023-30445-x.

[23] López-Blanco JR, Aliaga JI, Quintana-Ortí ES, Chacón P. IMODS: Internal coordinates normal mode analysis server. Nucleic Acids Res 2014;42. 10.1093/nar/gku339.

[24] Afshan G, Yaseen N, Ali SH, Khan AU. Immunoinformatics-Based development of a Multi-Epitope vaccine candidate targeting coinfection by Klebsiella pneumoniae and Acinetobacter baumannii. BMC Infect Dis 2025;25. 10.1186/s12879-025-11242-5.

[25] Rapin N, Lund O, Bernaschi M, Castiglione F. Computational immunology meets bioinformatics: The use of prediction tools for molecular binding in the simulation of the immune system. PLoS One 2010;5. 10.1371/journal.pone.0009862.

[26] Stolfi P, Castiglione F, Mastrostefano E, Di Biase I, Di Biase S, Palmieri G, et al. In-silico evaluation of adenoviral COVID-19 vaccination protocols: Assessment of immunological memory up to 6 months after the third dose. Front Immunol 2022;13. 10.3389/fimmu.2022.998262.

[27] Barazani O, Erdos T, Chowdhury R, Kaur G, Venketaraman V. New Advances in the Development and Design of Mycobacterium tuberculosis Vaccines: Construction and Validation of Multi-Epitope Vaccines for Tuberculosis Prevention. Biology (Basel) 2025;14. 10.3390/biology14040417.

[28] Granados-Tristán, A. L., Hernández-Luna, C., González-Escalante, L., Camacho-Moll, M., Silva-Ramírez, B., De León, M. B., & Peñuelas-Urquides, K. (2023). ESX-3 secretion system in Mycobacterium: An overview.. Biochimie. 10.1016/j.biochi.2023.10.013

[29] Sharma R, Rajput VS, Jamal S, Grover A, Grover S. An immunoinformatics approach to design a multi-epitope vaccine against Mycobacterium tuberculosis exploiting secreted exosome proteins. Sci Rep 2021;11. 10.1038/s41598-021-93266-w.

[30] Jiang F, Han Y, Liu Y, Xue Y, Cheng P, Xiao L, et al. A comprehensive approach to developing a multi-epitope vaccine against Mycobacterium tuberculosis: from in silico design to in vitro immunization evaluation. Front Immunol 2023;14. 10.3389/fimmu.2023.1280299.

[31] Mohammadipour S, Tavakkoli H, Fatemi SN, Sharifi A, Mahmoudi P. Designing a multi-epitope universal vaccine for concurrent infections of SARS-CoV-2 and influenza viruses using an immunoinformatics approach. BMC Infect Dis 2025;25. 10.1186/s12879-025-11066-3.

[32] Mansouri A, Yousef MS, Kowsar R, Miyamoto A. Homology Modeling, Molecular Dynamics Simulation, and Prediction of Bovine TLR2 Heterodimerization. Int J Mol Sci 2024;25. 10.3390/ijms25031496.

[33] Colleselli K, Stierschneider A, Wiesner C. An Update on Toll-like Receptor 2, Its Function and Dimerization in Pro- and Anti-Inflammatory Processes. Int J Mol Sci 2023;24. 10.3390/ijms241512464.

[34] Sahoo S, Lee HK, Shin D. An integrated structural and immunoinformatic approach to design a multi-epitope based vaccine against the foot-and-mouth disease virus. Sci Rep 2025;15. 10.1038/s41598-025-19826-6.

[35] Turner JS, O’Halloran JA, Kalaidina E, Kim W, Schmitz AJ, Zhou JQ, et al. SARS-CoV- 2 mRNA vaccines induce persistent human germinal centre responses. Nature 2021;596:109–13. 10.1038/s41586-021-03738-2.

